# Assessing the potential of vision language models for automated phenotyping of *Drosophila melanogaster*

**DOI:** 10.1101/2024.05.27.594652

**Authors:** Giulia Paci, Federico Nanni

**Affiliations:** Laboratory for Molecular Cell Biology, University College London, London, UK; Research Engineering Group, The Alan Turing Institute, London, UK

**Author notes:** The authors contributed equally to the paper.

## Abstract

Model organisms such as *Drosophila melanogaster* are extremely well suited to performing large-scale screens, which often require the assessment of phenotypes in a target tissue (e.g., wing and eye). Currently, the annotation of defects is either performed manually, which hinders throughput and reproducibility, or based on dedicated image analysis pipelines, which are tailored to detect only specific defects. Here, we assess the potential of Vision Language Models (VLMs) to automatically detect aberrant phenotypes in a dataset of *Drosophila* wings and provide their descriptions. We compare the performance of one the current most advanced multimodal models (GPT-4) with an open-source alternative (LLaVA). Via a thorough quantitative evaluation, we identify strong performances in the identification of aberrant wing phenotypes when providing the VLMs with just a single reference image. GPT-4 showed the best performance in terms of generating textual descriptions, being able to correctly describe complex wing phenotypes. We also provide practical advice on potential prompting strategies and highlight current limitations of these tools, especially around misclassification and generation of false information, which should be carefully taken into consideration if these tools are used as part of an image analysis pipeline.

## Introduction

*Drosophila melanogaster* has been used extensively as a model organism for large-scale genetic screens (1), drug screening for therapeutic applications (2) and toxicity studies (3).One of the most typical readouts of these assays is the presence of aberrant phenotypes in a target tissue, most often the adult fly eye and wing. However, this requires painstaking manual annotation of the observed phenotypes, which is labour-intensive and prone to subjectivity (for example with the use of arbitrary “scores” for phenotype severity). We focus here on the analysis of wing phenotypes, which are especially challenging as they can be extremely varied.

Currently, a few tailored image analysis tools are available for the analysis of *Drosophila* wing morphology. For example, Wings4 is a semi-automated tool that fits splines to the wing veins and enables the quantification of wing sizes and shapes (4). FijiWings is a set of Fiji (5) macros tailored for the quantification of wing size and trichome density (cell number) (6) MAPPER is a recently developed tool based on machine learning to segment and classify wing intervein and vein regions and quantify several geometrical and pattern-based features (7). While these tools can be extremely valuable to obtain quantitative measurements of wing shapes, their use as part of a screening pipeline is hindered by two limitations:

- they rely on segmentation of wing regions and/or fitting of specific wing structures (margin, veins), which limits their applicability to severely affected wings (e.g., blistered / truncated wings);
- using them to detect heterogeneous wing phenotypes would require the users to define a large set of “aberrant” conditions (wing shape descriptors, trichome density, vein lengths, etc.).

We reasoned that the recent advancements in Generative AI (8), with the development of Vision Language Models (VLMs) capable of handling both text and images as input prompts and to produce text as output (such as GPT-4 Vision (9) or Gemini (10)), could aid researchers in phenotyping tasks. VLMs could be used to classify wing images between normal (wild type, WT) or abnormal and, crucially, provide textual descriptions of the phenotypes observed. If VLMs are capable of performing such tasks, they could be used to inspect a large number of images and “flag” samples showing defects for follow-up. Approaches as the one described here are part of the current transition in applied machine learning from supervised learning strategies (11) to in-context learning (12), namely the ability of a model to learn to perform a task based on the prompt description (so called “prompt engineering” (13)), without any explicit updates to its parameters (i.e., without fine-tuning).

Nevertheless, it is now also widely known that large language models are highly prone to creating false information, producing so-called “hallucinations” (14, 15). Therefore, thorough and extensive evaluations of their capabilities and mistakes are necessary before deciding whether to employ them in any downstream task.

With such mindset, in this study we leverage a published database of annotated *Drosophila* adult wing phenotypes (16) and assess the performance of different VLMs and prompting strategies to provide an accurate detection of abnormal wings and a description of the observed phenotypes (see a graphical example in Figure 1). We decide to focus on one of the currently best-performing models (according to Bitton et al. (17)), GPT-4 Vision, and a small-scale quantized open source alternative, LLaVA (18), which can be run on a laptop^1^.

**Fig. 1.**
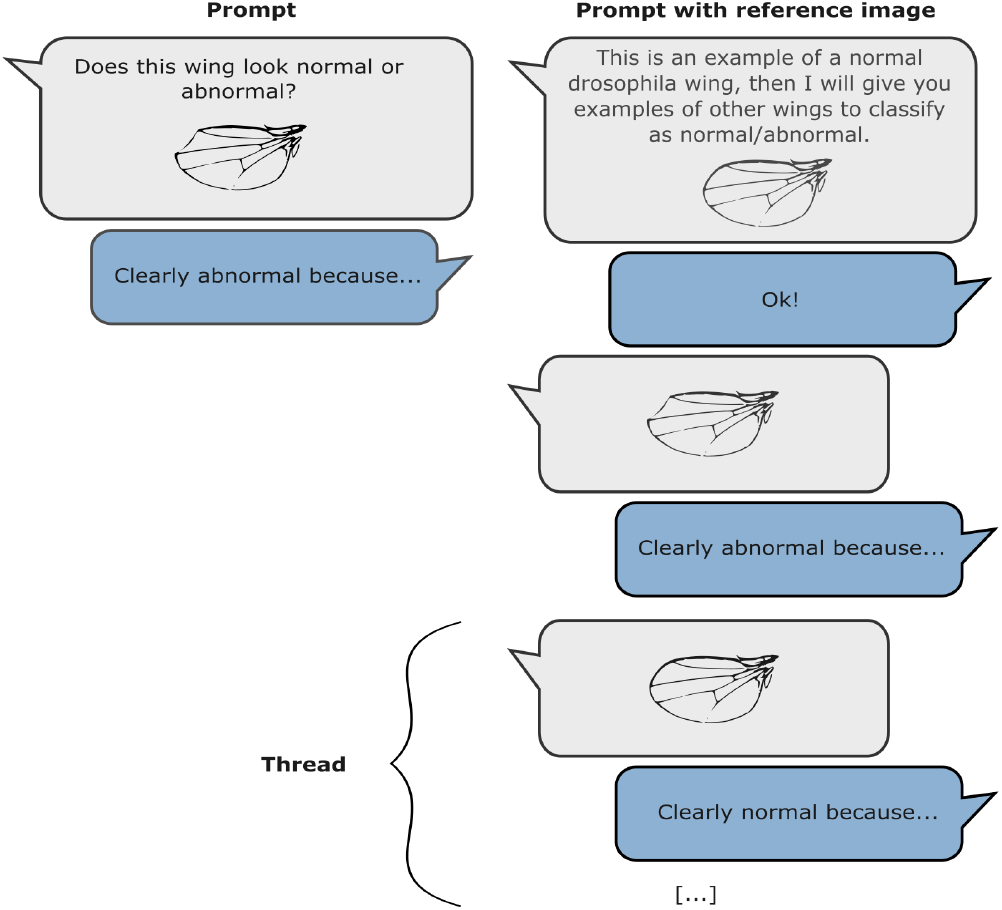
Schematic representation of the prompt strategies tested.

We assess the output generated by such models at three levels of granularity: first of all, we measure whether they are able to binary discriminate when presented with an image of a normal or abnormal wing. Secondly, by examining the descriptions generated, we verify whether the model correctly identifies the class of defect observed (e.g., a vein vs wing margin defect) and, finally, we ensure that the model does not produce any false information about the abnormality.

We also examine the performance of each model for the detection of different classes and degrees of severity of wing phenotypes. We conclude by discussing potential additional challenges when employing such methodology in practice^2^.

## Dataset

We employed wing images from a previously deposited dataset (16) where UAS-RNAi lines targeting 10,920 protein-coding genes were screened for phenotypes in the adult wing. We selected a subset of 80 images covering a wide range of phenotypes: ectopic wing veins, loss of wing veins, integrity of the wing margin, wing surface adhesion. Phenotypes ranged from weak ((w) annotation in the dataset, under the phenotype column) to severe ((s) annotation in the dataset). We also collected a set of 20 wings that were classified as phenotypically normal (“like-WT”) in the study (see overall statistics and examples in Table 1). Images, originally in PSD format, were converted to PNG as the language models adopted are not able to process PSD as an input file.

**Table 1.**
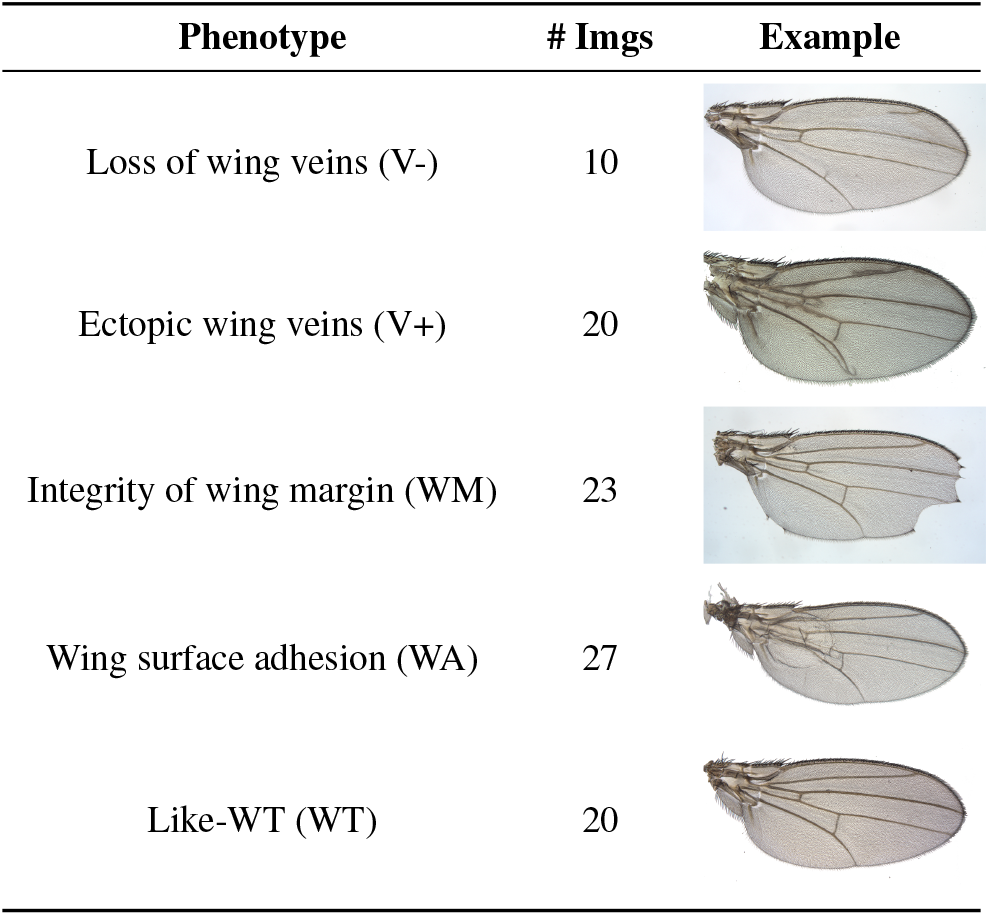
Dataset: phenotypes analyzed and example images.

| Phenotype | # Imgs | Example |
| --- | --- | --- |
| Loss of wing veins (V-) | 10 |  |
| Ectopic wing veins (V+) | 20 |  |
| Integrity of wing margin (WM) | 23 |  |
| Wing surface adhesion (WA) | 27 |  |
| Like-WT (WT) | 20 |  |

For tests where the VLM was provided with a reference wing image, we used a WT wing of Samarkand strain from Sonnenschein et al. (19) as reference, in order to make sure the VLMs would be able to generalise wing properties beyond the type of image taken^3^. The selected image (in TIF format) was cropped to remove scale bar and flipped to match orientation of the dataset used, then exported as PNG.

## Methods

We tested two Vision Language Models, GPT-4 and LLaVA in the following settings:

**GPT-4** (Turbo) was tested through the online chat interface^4^. In our experiments, we decided to use GPT-4 through the interface to facilitate the annotation process, given we had to compare a large number of generated textual descriptions with provided images. However, in a real application, we would strongly recommend relying on the OpenAI API^5^, which guarantees more control of the model (e.g., fixing specific parameters) and greatly speeds up the analysis.

**LLaVA** (version: 1.6, parameters: 7B, quantisation: 4-bit, size: 4.1 GB) was used through Ollama^6^, a simple tool for setting up an API to run language models locally on a personal computer, with default parameters. Note that the model we used is quantised. Quantisation reduces the number of bits required for each parameter, considerably diminishing the memory needed. However this has an impact on model’s accuracy (to know more see Dettmers and Zettlemoyer (20)).

### Experimental Setting

Large Language Models have shown incredible performance when presented with textual descriptions of a new task (12), without any annotated examples (so called “zero-shot” prompting (21)). To assess this in the context of *Drosophila* phenotyping, we compared VLMs performance by only providing a textual prompt describing the task (strategy named “p”) and by providing the textual prompt together with a single reference image of a normal *Drosophila* wing (strategy “p+i”)^7^. See the prompts used in Table 2.

**Table 2.** Prompt strategies used in this study.

| Prompt Strategy | Textual Content |
| --- | --- |
| Prompt (p) | I want help to identify drosophila wings that show any abnormalities. Any defect in the veins, wing margin or wing texture as well as blistering will count as abnormal. Does this wing look normal or abnormal? [add image to annotate] |
| Prompt with Reference Image (p+i, specifically for GPT-4) | I want help to identify drosophila wings that show any abnormalities. This is an example of a normal drosophila wing and I will give you examples of other wings to classify as normal/abnormal. Any defect in the veins, wing margin or wing texture as well as blistering will count as abnormal [add reference image] ... [model response, confirming it understood the task] ... [add image to annotate] |
| Prompt with Reference Image (p+i, specifically for LLaVA) | I want help to identify drosophila wings that show any abnormalities. Any defect in the veins, wing margin or wing texture as well as blistering will count as abnormal. In the image I'm providing you have on top an example of a normal drosophila wing and below a second example. Does this second wing look normal or abnormal compared to the first? |

To further test the ability of VLMs to leverage in-context information, we ran another series of experiments where we provided wing images to be classified one after the other as part of a thread, starting again with a reference image. We randomise image order, to make sure the model would not pick any obvious signal based on the sequence of images. While no feedback was given to the tool for each generated description, we wanted to understand whether performing the same task multiple times in a thread would help the VLM to more accurately identify phenotypes. We compared two different strategies, one in which all 100 images were given one by one in the same thread (from now on “p+i+100t”) and a second strategy where we provided the VLM images in threads of ten by ten, with the reference image shown at the beginning of each thread (from now on “p+i+10t”). Note that, since LLaVA can handle only one image at a time, we performed such experiments only with GPT-4.

### Evaluation Metrics

We employed several machine learning metrics to evaluate the effectiveness of VLMs in accurately discriminating normal and abnormal wing phenotypes. Note that, occasionally, the models provide an inconclusive output: we considered these as classification errors. Initially, we focused on the model accuracy, which quantifies the proportion of total correct predictions, providing a straightforward measure of the model overall performance in identifying wing anomalies.

To better understand the strengths and weaknesses of each method, we employed precision and recall. Precision measures the accuracy of positive predictions; essentially, the lower the number of false positive for each category, the higher the precision. Meanwhile, recall assesses the model’s ability to identify all relevant instances of a specific class, thereby for instance indicating the model’s sensitivity to detecting anomalies. The F1-Score, which harmonises precision and recall through their harmonic mean, was also reported to provide a balanced view of model efficacy. Further, we aggregated these findings using the Macro F1-Score, offering a comprehensive overview that equally considers the performance across both classes without being influenced by class imbalance (which in our case is high, as we have 80 images with defects and 20 controls).

Lastly, we examined the error rate, which reflects the proportion of all incorrect predictions to highlight specific areas where the model’s performance may lag, for instance in the tail of a long thread. These metrics were instrumental to underscore the limitations of current methodologies and identify potential areas for improvement in the automated analysis of phenotypes.

## Results

### Model comparison: GPT-4 and LLaVA

In Table 3, we compare the general performance of GPT-4 and LLaVA in distinguishing between images of wings with defects compared to control examples, in terms of precision, recall and F1-Score. We report these metrics for each class and also the overall model accuracy and Macro F1-Score, which is the average of the F1-Score for each class. These different metrics help understand where each model performs well in this general binary task.

**Table 3.** Results of model evaluations.

| Model | Accuracy | Aberrant phenotypes |  |  | WT phenotypes |  |  | Macro F1 |
| --- | --- | --- | --- | --- | --- | --- | --- | --- |
|  |  | Precision | Recall | F1 score | Precision | Recall | F1 score |  |
| GPT-4 (p) | 0.35 | **1.00** | 0.19 | 0.32 | 0.24 | **1.00** | 0.38 | 0.35 |
| GPT-4 (p+i) | **0.69** | 0.93 | 0.66 | 0.77 | **0.37** | 0.80 | **0.51** | **0.64** |
| LLaVA (p) | 0.37 | 0.77 | 0.30 | 0.43 | 0.19 | 0.65 | 0.29 | 0.36 |
| LLaVA (p+i) | 0.66 | 0.79 | **0.78** | **0.78** | 0.18 | 0.20 | 0.19 | 0.49 |
| Random baseline | 0.50 | 0.81 | 0.49 | 0.61 | 0.21 | 0.55 | 0.31 | 0.46 |

Both GPT-4 and LLaVA perform very poorly (below a reference baseline, which classifies images randomly) when prompted without a reference image (p strategy). This is particularly clear for GPT-4, which misses the majority of wings with defects (19% recall for aberrant phenotypes). We can take this as a reflection of the models poor “a priori knowledge” of how a normal *Drosophila* wing should look like. Interestingly, both GPT-4 and LLaVA sometimes provided answers including atypical terminology for the veins: *“[*…*] In a typical Drosophila wing, there are several key veins that should be present and unbroken: the costa, subcosta, radius, media, cubitus, anal, and the crossveins. [*…*]”. Drosophila* wing veins are typically divided into longitudinal veins (L1-5) and crossveins (ACV, PCV), while this terminology is more often used to describe conserved features of overall insect anatomy.

Given this starting point, the improvement in GPT-4 performance across most metrics when provided with just an individual reference image (p+i strategy) is impressive. In particular, an accuracy of nearly 70% and a F1 score of over 75% in detecting wings with defects show some clear potential for downstream applications. LLaVA (p+i) was also able to detect wings with aberrant phenotypes with high performance, however it greatly over-annotated wings as having defects, as can be seen by its very poor results on WT phenotypes (see Table 3).

From these results it is already clear that the task of automatic recognition of phenotypes currently cannot be completely automatised, but VLMs could support researchers and speed up the initial screening phase (for instance, LLaVA took less than 3 seconds to examine and generate a description of each image pair). This is especially true in a prototyping phase: testing the capabilities of such approaches takes at most a few lines of Python code (or can be done through an online chat interface, like the case of GPT-4). This is very different compared to a traditional supervised machine learning classification process, which would require obtaining training data, setting up a pipeline and testing different algorithms before seeing the first results.

### Testing GPT-4 thread

As a second step, we decided to explore whether providing images to annotate as part of a thread (instead of single images independently) would help or hinder performance. We compared two strategies, either providing all images one by one as part of a single thread or providing images in 10 threads, each containing 10 images. Note that input images were randomized across phenotypes for all thread tests.

In Figure 2, we report a comparison of error-rates in chunks of 10 examples; these chunks correspond to the input of each thread in the p+i+10t approach. We compared the two thread approaches with the GPT-4 (p+i) approach, which was the one performing best in Table 3. As can be clearly seen, the error rate of prediction for the single thread approach (p+i+100t) greatly increases when provided with more than 50-60 examples, with the outputs indicating that the model had seemingly “forgotten” the task at hand. On the other hand, the 10 threads approach (p+i+10t) shows very good and consistent performance, very much in line with the p+i approach tested before. From this point on, we focus subsequent performance analyses on the p+i approaches (GPT-4 and LLaVA) as well as the 10 threads (p+i+10t), which were found to be the most promising strategies.

**Fig. 2.**
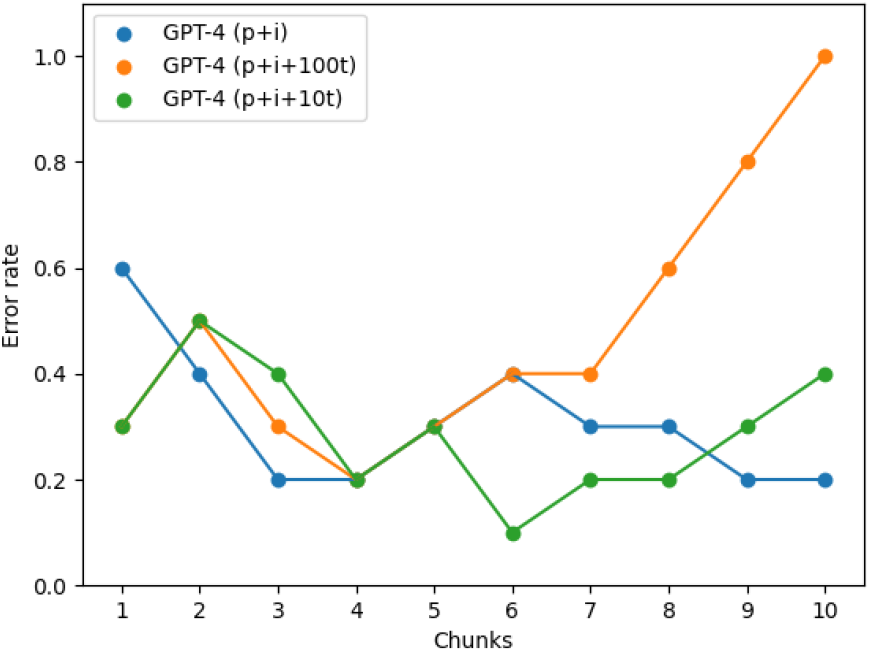
Error rate of the thread and the prompt+image strategies for GPT-4, in chunks of ten images.

### Performance by Phenotype

In Table 4 we further examine the models performance by breaking down their evaluation for the different phenotype classes. First of all, it is evident that LLaVA (p+i) over-classifies wings as having defects, as can be seen by its extremely low accuracy for control images (20%, compared with an accuracy of 80% and 85% of the GPT-4 based approaches). On the other hand, the GPT-4 approaches find it particularly difficult to recognise aberrant phenotypes related to wing veins.

**Table 4.** Accuracy by phenotype class.

| Phenotype | GPT-4 (p+i) | LLaVA (p+i) | GPT-4 (p+i+10t) |
| --- | --- | --- | --- |
| V+ | 0.45 | **0.80** | 0.55 |
| V- | 0.30 | **0.80** | 0.40 |
| WM | 0.74 | 0.65 | **0.78** |
| WA | **0.89** | 0.85 | 0.78 |
| WT | 0.80 | 0.20 | **0.85** |

We wondered whether the poor performance for some models and phenotype combinations could be due to the strength of the phenotype to be analysed (e.g., too weak), and sought to test this. In Figure 3, we report the accuracy of the models for the binary classification task, split by the strength of the phenotype: strong or weak, as reported in the published dataset. Phenotypes without specific strength annotation are considered of normal intensity. As can be noticed, both GPT-based strategies perform similarly, with very high accuracy (over 80%) for images with strong phenotypes and a drastic decrease for images with weak phenotypes (reaching around 10% for the p+i+10t approach). To further stress the fact that LLaVA’s outputs are unspecific, the performance for the three groups are counter-intuitively very similar with each other, and especially high when examining weak phenotypes.

**Fig. 3.**
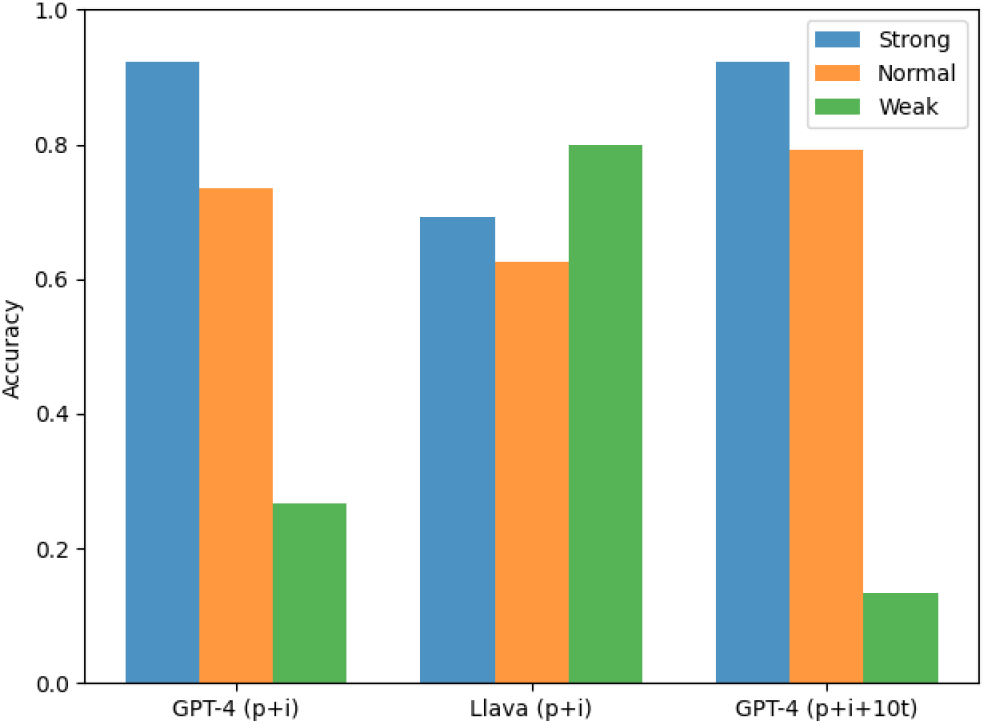
Accuracy of defect classification by phenotype strength.

### Evaluation of output descriptions

In Table 3, we have assessed whether VLMs are able to distinguish in a coarse-grained way between wings with aberrant phenotypes (defects) versus WT. However, the power of VLMs in comparison with traditional supervised classification pipelines lies in the textual description that they generate. Therefore, we have further assessed the quality of the descriptions provided by the models with two additional checks:

- whether the model correctly identifies in the description the present phenotype class (for example, veins are mentioned for a V+ or V-wing image);
- whether it does not produce any false information about the phenotype.

As can be seen in Figure 4, LLaVA is the overall best performing method in detecting wings with defects (with a tendency of over-classifying wings with having defects, as already discussed in relation to Table 4). However, when examining whether the descriptions do in fact mention the correct defect, we see a decrease in performance of around 30%. This is due to the fact that most of LLaVA’s generated descriptions contain vague and false information. This is further highlighted by the fact that less than 10% of images with defects have been presented with a description that does not contain any wrong information. This is clearly alarming for any real application and so it is important to underline that currently the potential of using a small VLM like LLaVA might only reside in the initial filtering of wings with and without defects, but they are very prone to hallucinations and may provide textual descriptions of limited usefulness.

**Fig. 4.**
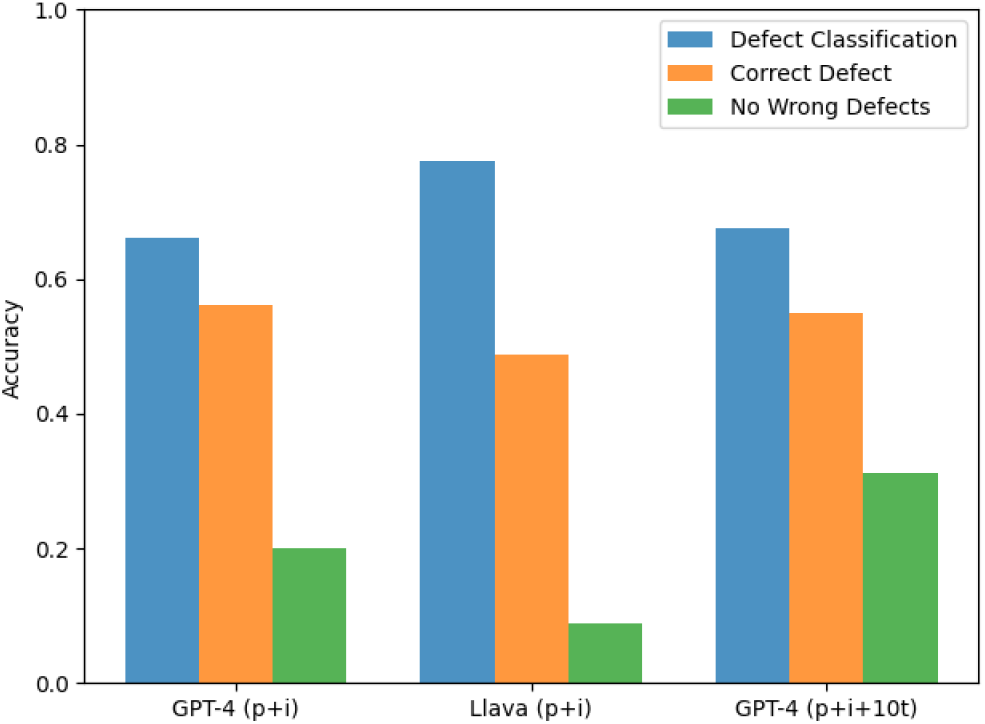
Comparison of output description quality.

Results for GPT-4 show a less drastic decrease in performance when compared to LLaVA, both in terms of mentioning the correct defect and avoiding hallucinations. The advantages of the p+i+10t approach are evident in this setting, were in 55% of the images of wings with defects the correct defect is mentioned in the description and 30% of descriptions do not contain any wrong information.

In Table 5 we provide some examples of model outputs that passed both checks, which highlight the potential of these models to annotate a wide range of phenotypes with varying degrees of severity. In Table 6 we further provide examples of model “hallucinations” and incorrect outputs.

**Table 5.**
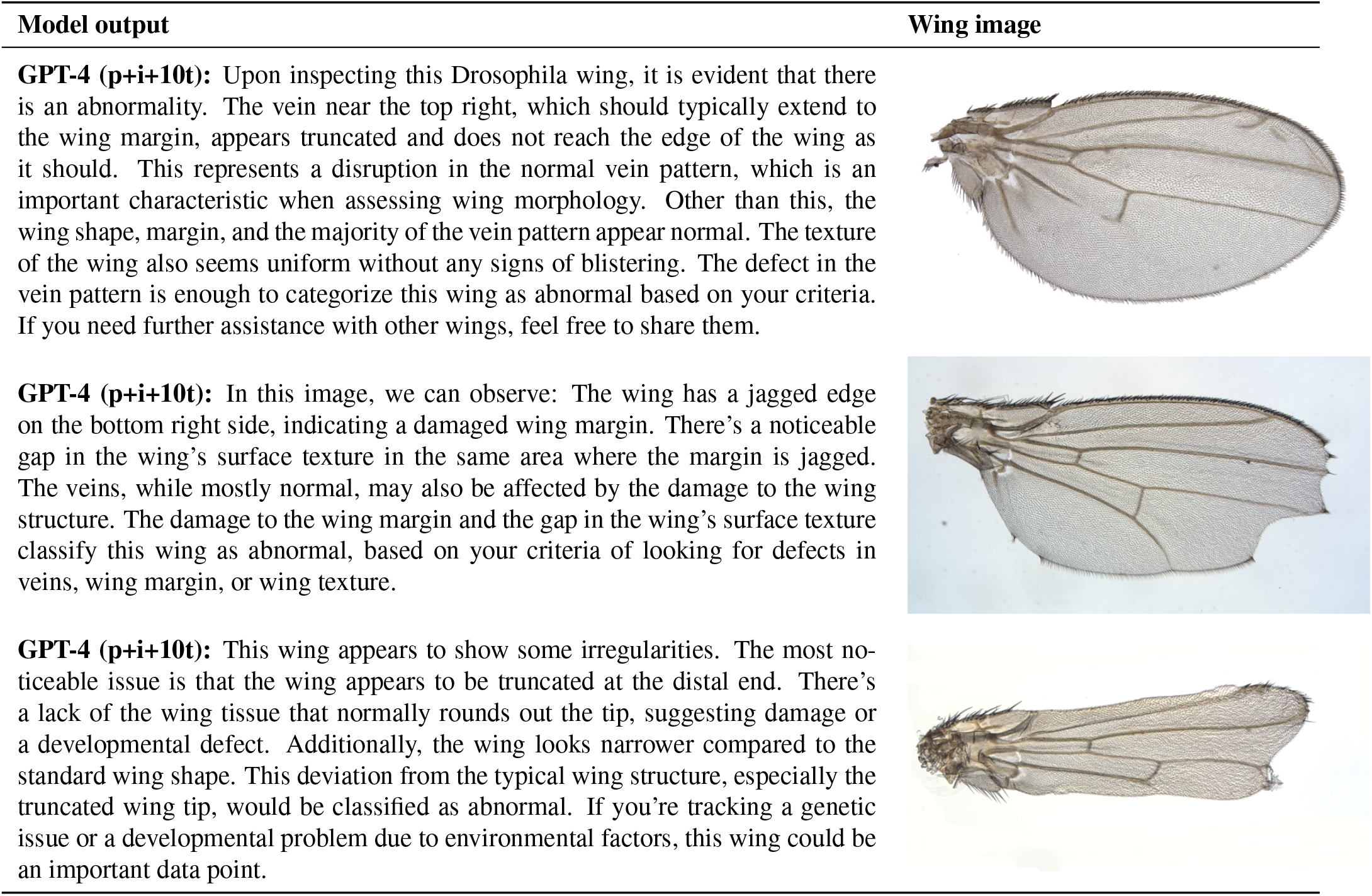
Examples of correct model outputs.

| Model output | Wing image |
| --- | --- |
| **GPT-4 (p+i+10t):** Upon inspecting this *Drosophila* wing, it is evident that there is an abnormality. The vein near the top right, which should typically extend to the wing margin, appears truncated and does not reach the edge of the wing as it should. This represents a disruption in the normal vein pattern, which is an important characteristic when assessing wing morphology. Other than this, the wing shape, margin, and the majority of the vein pattern appear normal. The texture of the wing also seems uniform without any signs of blistering. The defect in the vein pattern is enough to categorize this wing as abnormal based on your criteria. If you need further assistance with other wings, feel free to share them. |  |
| **GPT-4 (p+i+10t):** In this image, we can observe: The wing has a jagged edge on the bottom right side, indicating a damaged wing margin. There's a noticeable gap in the wing's surface texture in the same area where the margin is jagged. The veins, while mostly normal, may also be affected by the damage to the wing structure. The damage to the wing margin and the gap in the wing's surface texture classify this wing as abnormal, based on your criteria of looking for defects in veins, wing margin, or wing texture. |  |
| **GPT-4 (p+i+10t):** This wing appears to show some irregularities. The most noticeable issue is that the wing appears to be truncated at the distal end. There's a lack of the wing tissue that normally rounds out the tip, suggesting damage or a developmental defect. Additionally, the wing looks narrower compared to the standard wing shape. This deviation from the typical wing structure, especially the truncated wing tip, would be classified as abnormal. If you're tracking a genetic issue or a developmental problem due to environmental factors, this wing could be an important data point. |  |

**Table 6.**
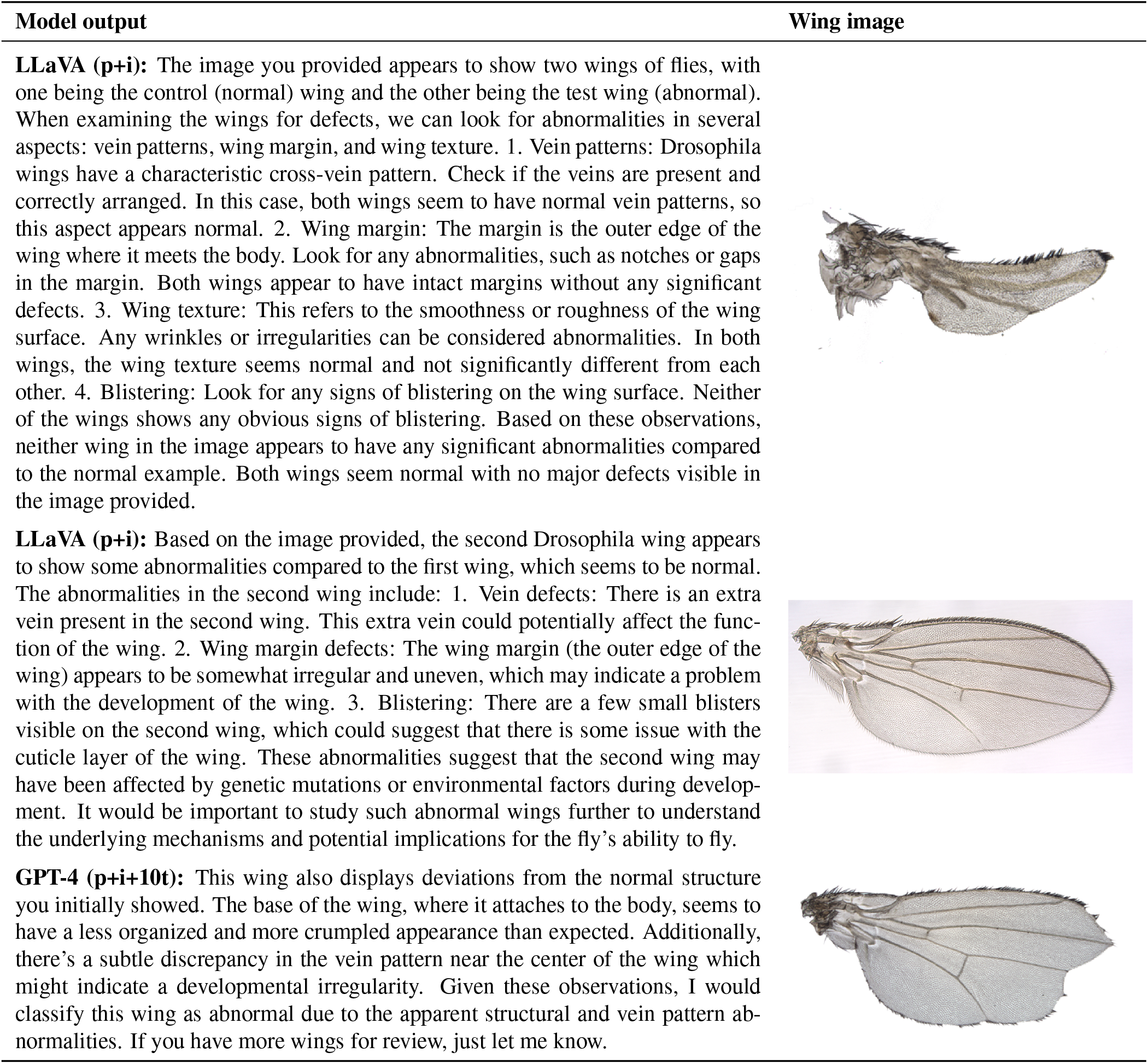
Examples of incorrect model outputs.

| Model output | Wing image |
| --- | --- |
| **LLaVA (p+i):** The image you provided appears to show two wings of flies, with one being the control (normal) wing and the other being the test wing (abnormal). When examining the wings for defects, we can look for abnormalities in several aspects: vein patterns, wing margin, and wing texture. 1. Vein patterns: *Drosophila* wings have a characteristic cross-vein pattern. Check if the veins are present and correctly arranged. In this case, both wings seem to have normal vein patterns, so this aspect appears normal. 2. Wing margin: The margin is the outer edge of the wing where it meets the body. Look for any abnormalities, such as notches or gaps in the margin. Both wings appear to have intact margins without any significant defects. 3. Wing texture: This refers to the smoothness or roughness of the wing surface. Any wrinkles or irregularities can be considered abnormalities. In both wings, the wing texture seems normal and not significantly different from each other. 4. Blistering: Look for any signs of blistering on the wing surface. Neither of the wings shows any obvious signs of blistering. Based on these observations, neither wing in the image appears to have any significant abnormalities compared to the normal example. Both wings seem normal with no major defects visible in the image provided. |  |
| **LLaVA (p+i):** Based on the image provided, the second *Drosophila* wing appears to show some abnormalities compared to the first wing, which seems to be normal. The abnormalities in the second wing include: 1. Vein defects: There is an extra vein present in the second wing. This extra vein could potentially affect the function of the wing. 2. Wing margin defects: The wing margin (the outer edge of the wing) appears to be somewhat irregular and uneven, which may indicate a problem with the development of the wing. 3. Blistering: There are a few small blisters visible on the second wing, which could suggest that there is some issue with the cuticle layer of the wing. These abnormalities suggest that the second wing may have been affected by genetic mutations or environmental factors during development. It would be important to study such abnormal wings further to understand the underlying mechanisms and potential implications for the fly's ability to fly. |  |
| **GPT-4 (p+i+10t):** This wing also displays deviations from the normal structure you initially showed. The base of the wing, where it attaches to the body, seems to have a less organized and more crumpled appearance than expected. Additionally, there's a subtle discrepancy in the vein pattern near the center of the wing which might indicate a developmental irregularity. Given these observations, I would classify this wing as abnormal due to the apparent structural and vein pattern abnormalities. If you have more wings for review, just let me know. |  |

## Discussion

In this study we have assessed the potential of vision language models for the classification and annotation of *Drosophila* phenotypes. We decided to focus this proof-of-principle test on wing phenotypes, as they display a remarkable range of variability, however the same strategies could be adopted for other organs. For example, they could be readily applied to detect *Drosophila* eye phenotypes, another typical target tissue used in genetic and drug screening.

Our study identifies current limitations and potential pitfalls of integrating VLMs in automated image analysis pipelines. LLaVA, a small scale open-source model, did not perform as well at GPT-4 in our tests. It has a tendency to over-classify wings as abnormal (only 20% accuracy for WT wings) and does not provide accurate descriptions of the phenotypes, often including irrelevant or wrong information. Despite these issues, LLaVA could still be valuable in specific cases, for example for the initial “flagging” of abnormal phenotypes in cases where false positives are not an issue. We found that GPT-4 performance when queried in a long thread dramatically decreased after 50-60 images, so this should be taken into account when designing the prompt strategy to use. Providing the data as “chunks” (e.g., 10 images) together with the reference image gave the best performance in our hands. Nevertheless, it is clear from our analyses that these tools are still far away from fully automatising a phenotype screening process and human supervision is still needed to double-check the results. Furthermore, all models appear very insensitive to changes in wing shape and size: for applications where the potential phenotypes involve subtle variations, adhoc pipelines like MAPPER or FijiWings are a better choice.

Despite the current limitations, VLMs also clearly show a lot of potential for future applications in this area. With a high accuracy of 80-90% for normal to strong phenotypes and overall output of 30% accurate descriptive outputs (phenotypes correctly identified, with no wrong information added), the best performing strategy (p+i+10t) could already be a valuable addition to image analysis pipelines. As can be seen in the examples in Table 5, this model can provide accurate descriptions of complex phenotypes, which go beyond the binary classification (normal/abnormal) and even the defect “classes” (veins, wing margin, etc.). Additionally, by providing a reference image, we only began exploring the potential of in-context learning and already obtained great improvements over the initial results. We therefore encourage researchers to explore other few-shot learning strategies (12), for instance offering reference images for all target defects to be detected or providing feedback to the model on the generated outputs.

In this proof of concept, we leveraged an existing dataset of annotated phenotypes based on images of dissected and mounted *Drosophila* wings. The images were all very clear and consistent (size, orientation), which simplified the task. However, we envision that VLMs could be integrated in pipelines capable of handling more complex scenarios, such as the detection of defects in images of whole *Drosophila* adults. For example, image segmentation approaches based on Segment Anything (22) could be used to identify target organ/s (eye, wings, legs) and then pass the segmented image to VLM-based downstream phenotyping systems. This would avoid the requirement of manually dissecting and mounting tissues, massively improving throughput.

In addition to this, large collections of mounted *Drosophila* wings (and other insects / tissues) already exist within databases, publications and digitised museum collections. VLMs could be extremely useful as part of pipelines to “mine” these images for a phenotype of interest and obtain information about the relevant genotype and conditions. These descriptions could then be integrated into existing databases such as FlyBase (23). In this study, we found that VLMs performance greatly increased by providing just an individual reference image, so they could readily be applied to different insect species and potentially used to discriminate them with minimal references needed.

To conclude, while visual language models are in their infancy, they already show potential for multiple applications in automated phenotyping studies. We encourage the community to carefully test them, being aware of their limitations and leveraging upon their strength to advance interdisciplinary research in this exciting new area between biology and artificial intelligence.

## Acknowledgements

We thank our colleagues Pablo Vicente Munuera, Courtney Lancaster and Ryan Chan for helpful comments and discussion. We also thank our little son, Leonardo, for being very patient while we were discussing *Drosophila* wing phenotypes during dinner.

## Footnotes

1 In our experiments, a MacBook Pro with M1 Pro and 32 GB of memory.

2 Materials for reproducing this study are available at: https://github.com/giuliapaci/VLM-Drosophila-Phenotyping

3 In initial experiments, we compared the performance of using this reference image and an image of a “like-WT” from Lopez-Varea et al. (16) and obtained similar performance.

4 Experiments were run between 19th of April 2024 and 5th of May 2024

5 https://platform.openai.com/docs/guides/vision

6 https://ollama.com/library/LLaVA

7 Note that, as LLaVA can only process one image at a time, we concatenated the reference image and the image to classify as a single image to make sure the model would correctly process the input.

